# Selective impairment of spatial recognition memory and reduced frontal corticothalamic spine density following adolescent alcohol consumption

**DOI:** 10.64898/2025.12.02.691762

**Authors:** Grace Qian, Hannah Jarrell, Faith Maxwell, Hernan Mejia-Gomez, Michael C. Salling

## Abstract

Heavy alcohol use is common during adolescence and is associated with increased risk of alcohol use disorder and prevalence of residual cognitive deficits, especially in behaviors associated with the latently developing prefrontal cortex (PFC). A major need for advancing our understanding of this relationship is replicable and accessible preclinical behavioral batteries that can be used to disassociate the effects of adolescent alcohol on select PFC circuits. Electrophysiological evidence implicates projections from the PFC to mediodorsal thalamus (PFC-MdT) as being uniquely impacted by adolescent intermittent alcohol consumption in mice. The present study aims to evaluate if voluntary consumption of alcohol during adolescence impacts anxiety or PFC-associated spatial and recognition memory and if they are associated with morphological changes to the PFC-MdT circuit. Our results indicate that compared to water-only controls, male and female mice that voluntarily consumed alcohol during adolescence demonstrate performance deficits in the object-in-place recognition task, without affecting the novel object recognition task, Y-maze alternation, anxiety or locomotion, an outcome consistent with PFC and MdT dysfunction. Morphological assessment of PFC neurons from male and female mice following behavioral tasks found a reduction in spine density in basal and apical, but not oblique dendrites of PFC-MdT neurons. Collectively, these results implicate the PFC to MdT circuit integrity as a potential locus of the effects of adolescent alcohol on spatial recognition memory.

## Introduction

Heavy alcohol consumption during adolescence alters the normal trajectory of brain development ^1^. The prefrontal cortex (PFC) is particularly vulnerable to the effects of adolescent alcohol consumption, as it is one of the last brain regions to achieve grey and white matter stabilization during adolescence due to extended synaptic pruning and myelination ^2^. Studies have shown that heavy alcohol drinking during adolescence significantly disrupts PFC grey and white matter trajectories and alters PFC activity ^3, 4^. Functionally, heavy alcohol consumption during adolescence produces deficits in behaviors associated with the PFC, including spatial working memory and response inhibition, as observed in clinical studies and preclinical rodent models ^5, 6^. Morphological assessment of prefrontal neurons in rodents indicates that adolescent alcohol exposure can reduce prefrontal grey matter as well as alter functional connectivity between the medial PFC and connected brain regions including striatal and thalamic outputs, supporting the validity of this line of preclinical investigation ^7, 8^. These structural and functional findings offer a candidate mechanism for adolescent alcohol’s associated effects on PFC-dependent cognitive control.

The PFC mediates executive function by integrating information from multiple cortical and subcortical regions ^9^. PFC function is largely tied to its interactions with the mediodorsal thalamus (MdT), which is the primary thalamic nuclei partner of the PFC with which it shares dense reciprocal connections ^10^. Studies in rodents have shown that damage to the MdT produces similar cognitive deficits to those seen after damage to the PFC, indicating that the MdT is a key node in maintaining PFC function including higher-order cognition^11^. Medial PFC (mPFC) to MdT circuitry have been shown to play a critical role in working memory, cognitive strategy, and choice selection in rodents ^12, 13^.

Disruptions to these circuits may contribute to the working memory deficits and impulsivity observed after chronic adolescent binge drinking ^14^. We have shown that intermittent access to alcohol (IA EtOH) during adolescence disrupts the acquisition of a working memory task assessed using a delayed non-match to sample T-maze task in addition to long-term effects on pyramidal neuron intrinsic excitability measured using whole-cell patch clamp electrophysiology ^6^. In follow-up experiments, PFC subpopulations were targeted using a retrograde viral approach, and male and female mice showed a reduction in glutamatergic transmission selectively in PFC^MdT^ pyramidal neurons, but not those projecting to the contralateral PFC ^15^. Therefore, the PFC^MdT^ circuit appears to be uniquely vulnerable to the effects of heavy alcohol consumption, particularly when initiated in early adolescence, and is likely a key contributor to cognitive dysfunction following adolescent alcohol exposure.

The medial PFC and MdT subregions have been functionally disassociated from hippocampal and rhinal subregions that regulate spatial and episodic encoding and instead regulate maintenance of memory traces and choice selection ^16^. Mice with bilateral lesions in the MdT or mPFC display deficits in an object-in-place associative recognition task, but not in a single-item novel object recognition task ^17^. Similarly, disconnection of the MdT and mPFC impaired performance on the object-in-place recognition task, but not the novel object recognition task, further implicating their shared circuitry^17^, thus providing a behavioral readout to disassociate PFC^MdT^ function from behaviors more associated with hippocampal and perirhinal lesions ^18, 19^. While previous studies using alcohol exposure models during adolescence have observed deficits on spatial and recognition memory, it is unclear how voluntary alcohol consumption during adolescence impacts performance on Y-maze and variations of object recognition tasks^20–22^. The present study was designed to investigate our hypothesis that adolescent binge drinking would lead to a selective effect on associative recognition memory and that PFC^MdT^ neurons would demonstrate morphological differences following adolescent alcohol consumption. Our results confirmed this initial hypothesis as we observed selective performance deficits on the mPFC- and MdT-dependent object-in-place associative recognition memory task and a reduction in spine density of PFC^MdT^ neurons following intermittent access to alcohol (IA EtOH) alcohol in male and female mice. Our results provide new insights into the vulnerability of the PFC^MdT^ circuit to adolescent alcohol consumption.

## Methods

### Subjects

All animal studies were performed according to the protocol approved by the Louisiana State University Health New Orleans Institutional Animal Care and Use Committee in accordance with the NIH Guide for the Care and Use of Laboratory Animals ^23^. Male (n = 17) and female (n = 19) C57BL/6J mice (stock# 000664, The Jackson Laboratory) between post-natal day (PND) 27-70 were used in this study and were randomly assigned to IA EtOH or control groups. Upon arrival in the laboratory, at ∼PND 21, mice were group housed under a reverse light/dark cycle (light phase: 22:00 to 10:00) with ad libitum access to food (autoclaved Teklad 2019S mouse chow) and water. Following surgical procedures, mice were individually housed, handled daily, and given nestlets for environmental enrichment for the remainder of the experiment. A timeline of experimental procedures is shown in **Figure 1A**.

**Figure 1:**
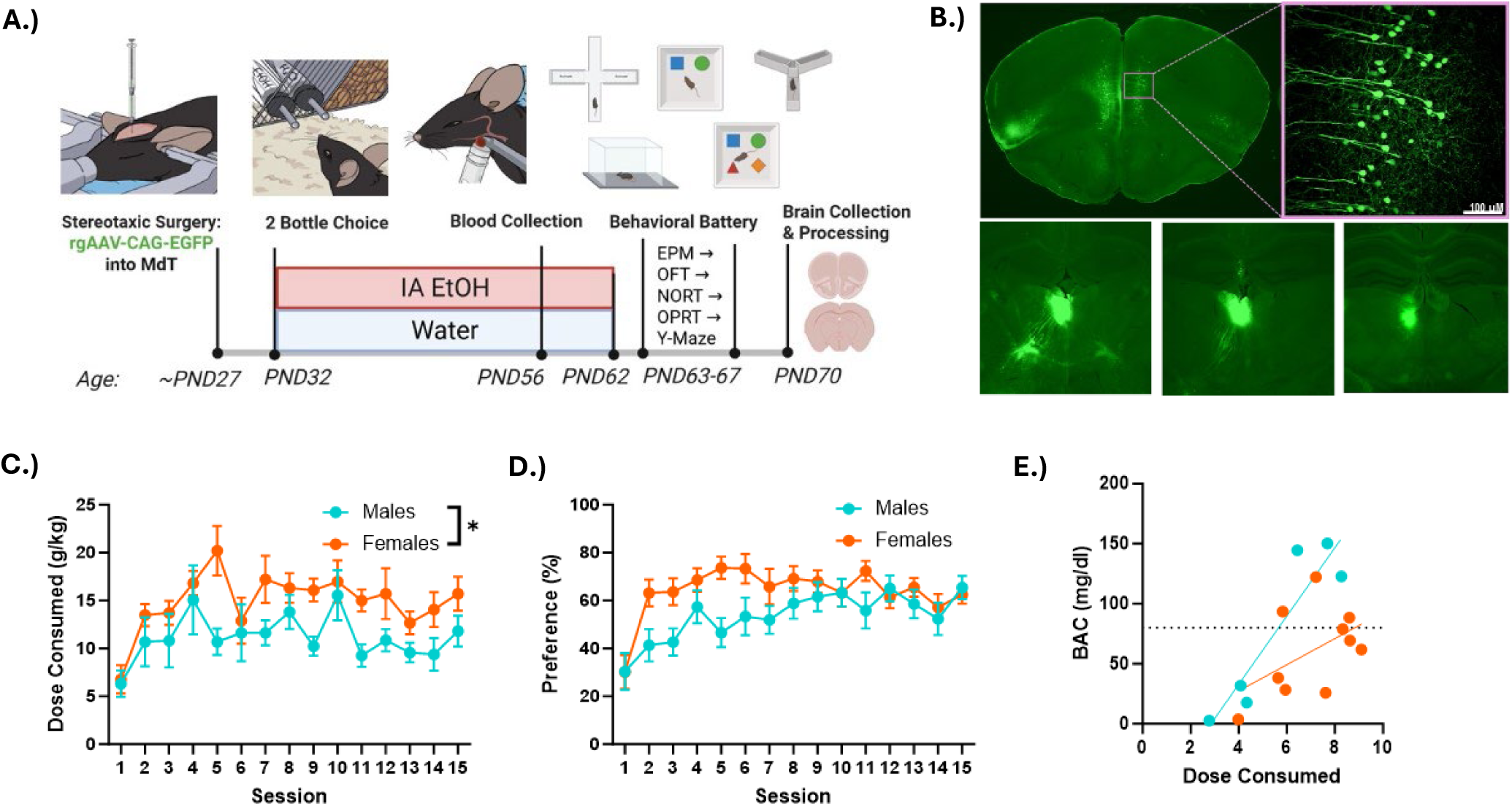
Experimental timeline and adolescent IA ETOH procedure: **A.)** Male and female mice underwent surgical procedures to infuse AAVrg-CAG-GFP into the MdT at PND 27 and began IA ETOH or water only consumption at ∼PND 32. All mice were placed on water only at ∼PND 62. The behavioral battery followed one day after last alcohol session and consisted of elevated plus maze (EPM), open field task (OFT), novel object recognition (NORT), object in place recognition task (OPRT), and Y-maze alternation over 5 days. Mouse brains were collected 8 days after last day of alcohol for dendritic spine analyses. **B.)** Top, left: representative images of GFP expression across the medial PFC and morphology of GFP expressing neurons (inset, top right). Bottom left to right: images of the range of injection sites included in dendritic spine analyses from the most anterior (-1.06 mm, from bregma), our target A-P coordinate (-1.2 mm), and most posterior injection site (-1.7 mm). **D.)** Under the IA EtOH procedure, male and female mice both escalated their dose consumed (g/kg), however female mice consumed more alcohol than male mice, p < 0.05. **E.)** Male and female mice demonstrated preference for the alcohol solution but did not statistically differ in their preference for the alcohol bottle. **F.)** 6 hours into the 13^th^ alcohol session (∼PND 58), a subset of mice reached blood alcohol concentrations (BACs) at or near binge levels (y = 80 mg/dl), however, under 24-hour access conditions, individual sampling at peak alcohol consumption time points is highly unlikely. Dose consumed was positively correlated with BACs in male, p < 0.001, but not female mice, p = 0.1, * p < 0.05. **1A:** Created in BioRender. Salling, M. (2025) https://BioRender.com/vvrapcu

### Surgical procedures

To enable visualization of PFC^MdT^ neurons, mice at PND 26-28 were anesthetized with continuous isoflurane (4% induction, 1.5% continuously thereafter), placed in the stereotaxic instrument (Kopf Instruments, Tujunga, CA) with temperature maintained using a heating pad (T/Pump CD2X). Following establishment of a surgical plane, mice received a subcutaneous (sq) injection of bupivacaine along the incision site, 0.5% ml sterile saline sq injection at the hindleg, a scalpel incision along the rostro-caudal axis of the head, followed by a cranial hole drilled over the infusion site. Mice next received an infusion of 200 nl virus (AAVrg-CAG-GFP, Addgene viral prep # 37825-AAVrg, 1x10^12^ vg/ml) unilaterally to the MdT (coordinates relative to bregma: AP -1.20 mm, ML ±0.35 mm, DV -3.10 mm) using a small gauge syringe (Hamilton Neuros 33g blunt tip) (**Figure 1A**). The viral injection side (right or left hemisphere) was counterbalanced within each sex and alcohol group. Mice were treated post-surgically with the analgesic Meloxicam SR, monitored for healthy recovery, and then individually housed thereafter to recover for at least 3 days prior to initiating the two-bottle choice drinking procedure and 4 days prior to their first alcohol session.

### Two-bottle choice intermittent alcohol consumption

To establish voluntary alcohol consumption, experimental mice were provided with intermittent access to alcohol (IA EtOH) through the two-bottle choice drinking protocol during adolescence (∼PND 32-62) similar to previous procedures ^6^. Individually housed mice were provided with two autoclaved 50 mL Nalgene bottles with double ball-bearing sippers containing either water or alcohol (15% v/v diluted from 95% ethanol, Decon Laboratories) (**Figure 1A**). Control mice were given two bottles of water, while mice from the IA EtOH group were given one bottle of alcohol and one bottle of water every other day and two bottles of water on intervening days. Bottles and mice were weighed daily, the position of the alcohol bottle (right/left) was alternated every other session, and a vacant drip cage was used to determine incidental fluid loss. Experimental mice were given a total of 15 alcohol consumption sessions over the 30 days. Alcohol dose consumed (g/kg) and alcohol preference (%) were calculated for each mouse after each session.

On the 13th alcohol access day, blood was collected from the submandibular vein at 6 hours into the dark cycle, and blood alcohol levels were measured using an Analox AM1 Analyzer. After the alcohol bottle was removed following the 15th alcohol session, mice began behavioral testing the following day.

### Behavioral Procedures

Following IA EtOH, mice underwent a battery of behavioral tasks over five consecutive days to assess the impact of adolescent alcohol consumption. Mice were tested on the elevated plus maze (EPM) and open field task on day 1, the novel object recognition task (NORT) on day 2, the object-in-place recognition task (OPRT) on day 3, and the Y-maze on day 4 (**Figure 1A**). All behavioral testing was conducted within the first 8 hours of the dark cycle. For all behavioral experiments, mazes and arenas were cleaned with Clidox S disinfectant and dried before each trial to remove odor cues and mouse waste. An infrared enabled camera (USB 2.0 Webcam, Microsoft) was secured above the apparatus and was used to record mouse activity during each trial. Videos were recorded and analyzed using open-source Bonsai software and location scored by blinded experimenters. Videos without complete recordings were excluded from the analyses. **Elevated Plus Maze (EPM)**: The EPM apparatus consisted of two open arms and two closed arms arranged perpendicularly to each other, with a center area, and was raised 50 cm from the floor (**Figure 2A**).

**Figure 2:**
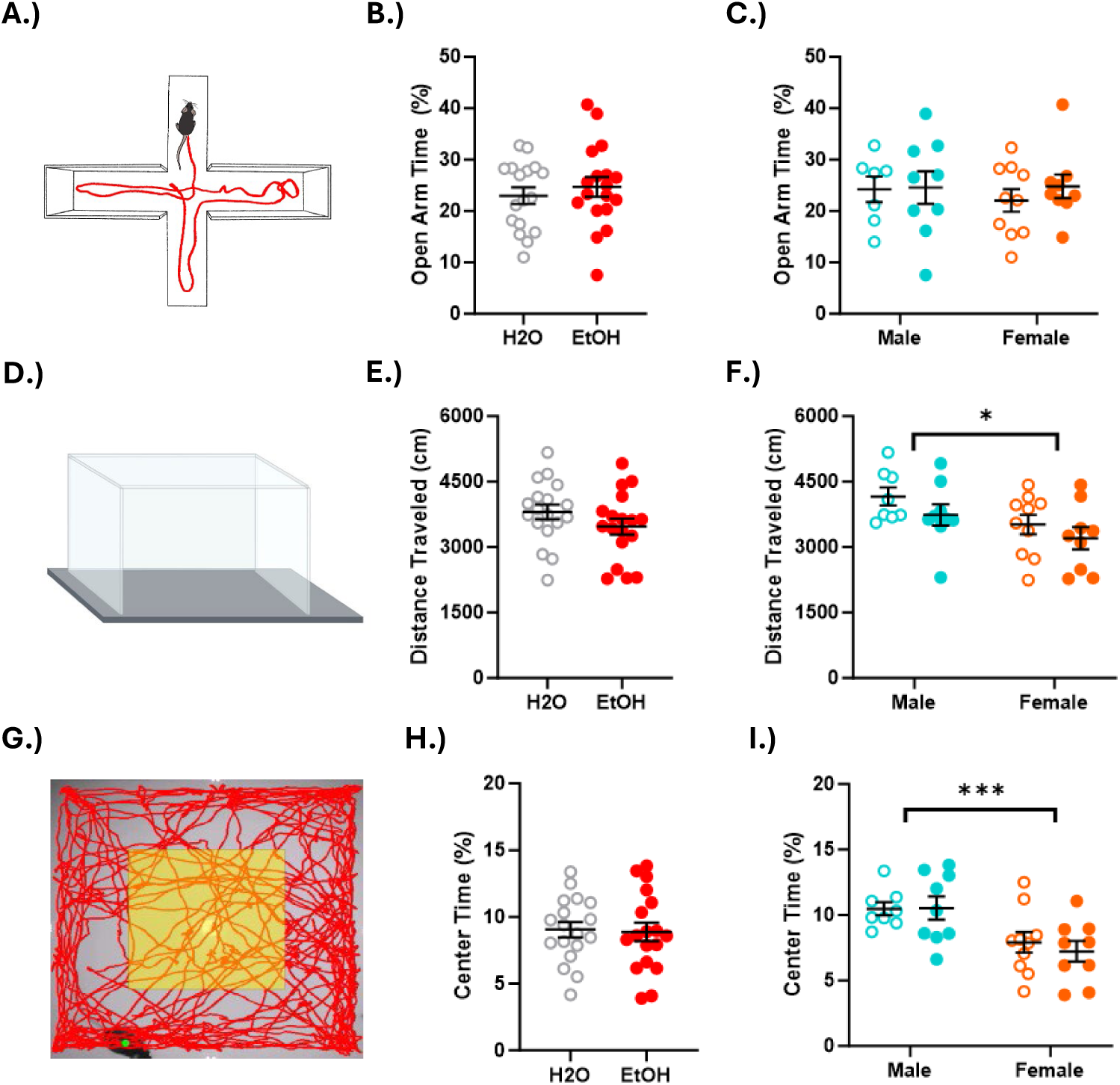
IA EtOH during adolescence did not affect anxiety or locomotor activity: **A.)** Illustration of EPM used to assess anxiety-like behavior. **B, C.)** Open arm time was not affected by adolescent IA EtOH or sex. **D.)** Illustration of open field task (OFT). **E, F.)** Spontaneous locomotor activity was not affected by adolescent IA EtOH, however, a main effect of sex was observed, p < 0.05. **G.)** Tracking recording of mouse demonstrating center arena used to measure center time. **H.)** Anxiety as assessed by percentage time in center of arena was not affected by adolescent IA EtOH. **I.)** A significant sex effect was observed, with female mice demonstrating increased anxiety-like behavior in the OFT, *** p < 0.005.

Each arm was 10 x 25 cm, and the two closed arms were enclosed by 20 cm high white plexiglass walls. The lighting within each arm of the maze was kept relatively uniform and dark (∼30 lux). Mice were taken from their home cage and placed in the center area of the maze facing the open arm opposite the experimenter who left the room following placement. Mouse location was tracked while exploring the maze for 5 minutes. An arm entry was defined as when all four of the mouse’s limbs crossed into the arm. Time spent on open arms was then expressed as a percentage of the total time spent on the maze. The number of entries into open arms and the percentage of time spent on open arms served as measurements of anxiety-like behavior. **Open Field Task (OFT):** The open field task was performed in an empty 40 x 40 x 40 cm black plexiglass box with a white floor, enclosed within a larger sound-attenuating behavioral chamber (Med Associates). Mice were taken from their home cage, placed in the middle of the arena, and recorded while moving freely throughout the arena for 8 minutes. The distance traveled and time spent in the center of the arena (30 x 30 x 30 cm) served as measurements of spontaneous locomotor activity and anxiety-like behavior, respectively. **Novel Object Recognition Task (NORT)**: The NORT took place the following day in the same arena as the open field task, which therefore served as a habituation phase prior to object-directed tasks. Each NORT trial consisted of three phases: sample, delay, and test phases (**Figure 3A**). In the sample phase, mice were taken from their home cage and placed in the arena containing two identical copies of the sample object, either two test tubes filled with colored gravel or two Lego towers, which were counterbalanced for sex and alcohol exposure, with mice location recorded while exploring the objects freely for 8 minutes. Mice were then returned to their home cage for the delay phase, lasting 5 minutes, during which time the box and objects were cleaned and dried to remove odor cues. In the test phase, mice were placed in the arena containing two objects, one identical to the sample object and one novel object, which was recorded while exploring the objects freely for 8 minutes. The number of novel interactions (NI), novel interaction time (NIT), familiar interactions (FI), and familiar interaction time (FIT) for each object during the sample and test phases were assessed from video recordings by an observer blind to experimental conditions. Object interaction was defined as the mouse pointing its nose towards the object within 2 cm of the object, including sniffing or touching the object. Time spent sitting on top of or climbing on the object was excluded from interaction time. The discrimination ratio (DR) of novel interaction time served as a measurement of single-item recognition memory and was calculated for the test phase using the following formula: (NIT – FIT)/(NIT + FIT). **Object-in-Place Recognition Task (OPRT)**: The OPRT was conducted in the same plexiglass box as the open field task and the NORT, however, 4 spatial cues were placed at the top of each wall. Similarly to the NORT, each OPRT trial consisted of sample, delay, and test phases. In the sample phase, mice were placed in the arena containing four unique objects located in the center of each of the four quadrants of the arena (**Figure 3D**). Mice were recorded while exploring the objects freely for 8 minutes using the same procedures as the NORT. In the test phase, mice were placed in the arena with the same four objects, except two objects switched positions while the other two objects remained in the same position as the sample phase. Following a 5-minute delay in their home cage, mice were next recorded while exploring the objects for 8 minutes. The objects that were moved in the test phase were counterbalanced across groups. The discrimination ratio was calculated for each trial like the NORT task except the novel stimulus was calculated as interactions with the two objects that had switched positions. **Y-maze**: The Y-maze consisted of three white, opaque, plexiglass arms (10 x 25 cm) at 120° angles from each other, labeled A, B, and C (**Figure 3G**). Each arm was enclosed by walls 20 cm high. Mice were brought into the room and placed in the center area of the maze. Mice were allowed to move freely through the maze and were recorded for 5 minutes. Each arm entry and the number of correct alternations were recorded manually from video recordings and scored by an observer blind to experimental conditions. An arm entry was recorded when all four of the mouse’s limbs were within an arm. A correct spontaneous alternation (SA) was defined as the mouse entering all three arms consecutively (e.g. B-C-A). The percentage of correct alternations served as a measurement of short-term spatial working memory and was calculated using the following formula: SA% = (SA/total alternations -2) x 100.

**Figure 3:**
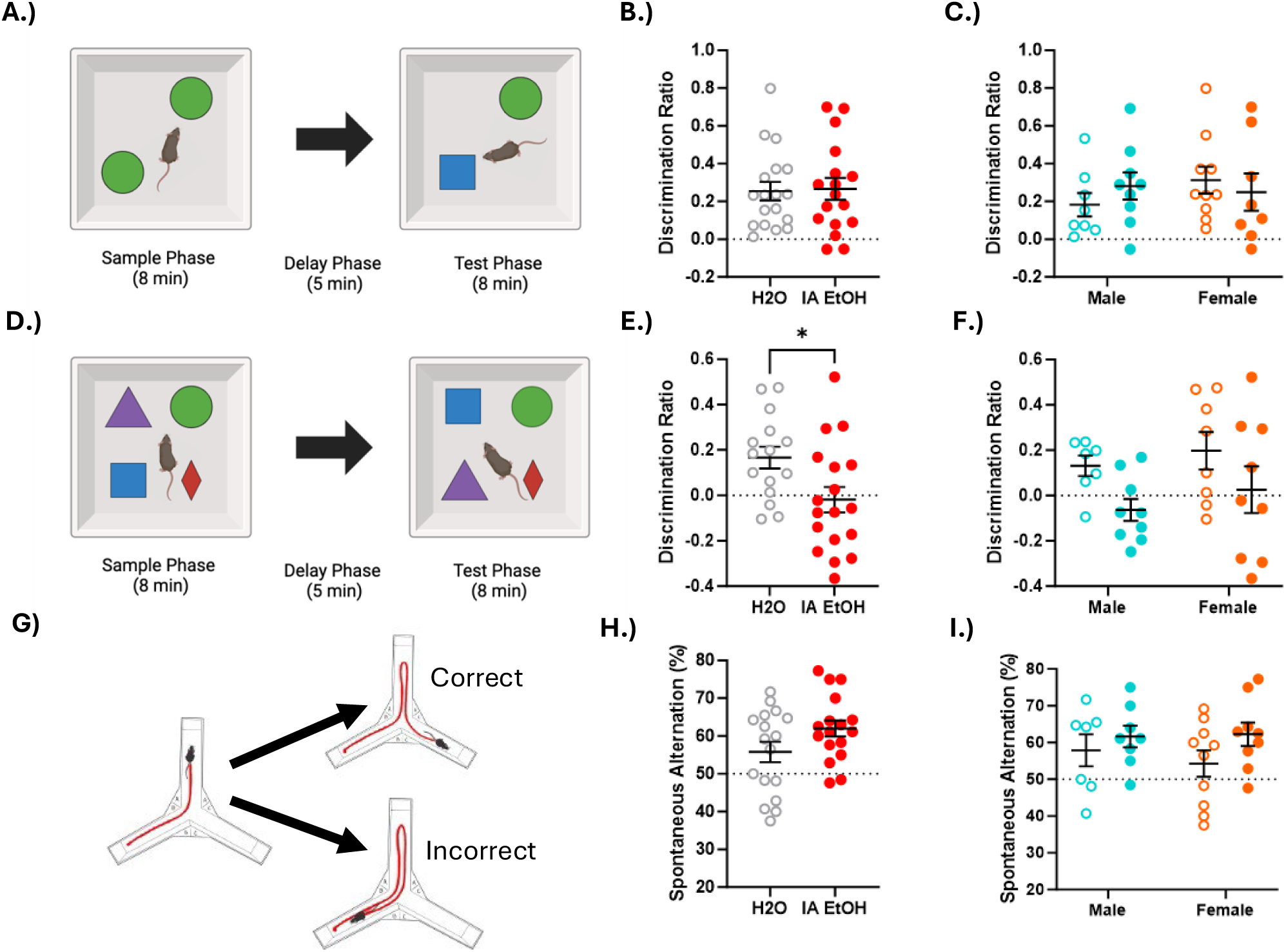
Assessment of cognitive function following adolescent IA EtOH selectively reduced associative recognition memory and not object or working memory: **A.)** Illustation of NORT, where mice were provided access to two identical objects in sample phase and one familiar and one novel object in test phase. **B.)** No effect of IA EtOH on discrimination ratio was observed in pooled or **C.)** sex independent comparisons. **D.)** Illustration of OPRT, where mice were provided with 4 different objects in sample phase and the same objects in test phase, but with the location of 2 objects switched. **E.)** Adolescent IA EtOH mice had a significant reduction in discrimination ratio, **F.)** not affected by sex. **G.)** Illustration of Y-Maze spontaneous alternation task where correct alternation was alternating between each arm and incorrect alternation was returning to previous arm. Y-maze alternation was not affected by **H.)** IA EtOH or **I.)** sex, * p < 0.05.

### Immunohistochemistry

Three days following completion of behavioral tasks, mice were anesthetized with 4% isoflurane and transcardially perfused with 0.1 M phosphate-buffered saline (PBS) followed by 4% paraformaldehyde (PFA) diluted in PBS. Heads were removed, degloved, and fixed overnight in 4% PFA in PBS. Brains were then extracted from the skull and sectioned at 50 μm thickness using a vibratome (Leica VT1200S). Free-floating sections were washed with PBS and then PBS with 0.3% Triton X (PBST), then incubated in blocking buffer (5% v/v goat serum in PBST) for 1 hour at room temperature. Sections were then incubated in primary antibody (chicken polyclonal IgY anti-GFP, Abcam, catalog # ab13970) diluted 1:2000 in blocking buffer for two nights at 4°C. Sections were washed with PBST and incubated in secondary antibody (AlexaFluor-488 goat anti-chicken IgY H&L, Abcam, catalog # ab150169) for 2 hours at room temperature. Sections were then washed with PBST and PBS before being mounted on slides using ProLong Diamond Antifade Mountant (ThermoFisher, catalog #P36961) and cover slipped.

GFP fluorescence was examined to identify neurons resulting from retrograde transduction and cytoplasmic expression in the neuron. First, sections were tile imaged at 10x on a fluorescence microscope (BZX800L/BZ-X810, KEYENCE) to confirm injection sites were in the MdT (**Figure 1B**) using a mouse atlas ^24^. Representative images of a slice containing the mPFC at 2x magnification and a mPFC neuron at 60x magnification are shown in **Figure 1B**. GFP expression in the slice containing the MdT was examined for each mouse. 24 mice showed a pattern of GFP expression within the MdT of axon tracts entering the MdT. Mice with missed MdT injections or minimal GFP expression in the medial PFC (n = 7) were included in the behavioral data but excluded from the morphological analyses. In some cases, proper morphology was not collected typically due to neuronal GFP expression that was too crowded to select out dendritic segments (n = 2) and mice were not included in the final morphology dataset.

### Morphological Analyses

For spine density measurements, segments were imaged using a confocal microscope (Leica DMi8) with a 100x, 1.4 mm NA objective and 488 nm laser. Image resolution was 2048 x 2048 pixels without zoom. 3D images were collected using a z-step of 0.28. Dendrite segments were cropped to 50 μm for analyses and classified as basal, oblique, or apical subtypes. Basal dendrites branched directly from the soma. Oblique dendrites were near to the soma but branched from the apical trunk. Apical dendrites were those located in the apical tufts in layers 1 and 2 (**Figure 4A**).

**Figure 4:**
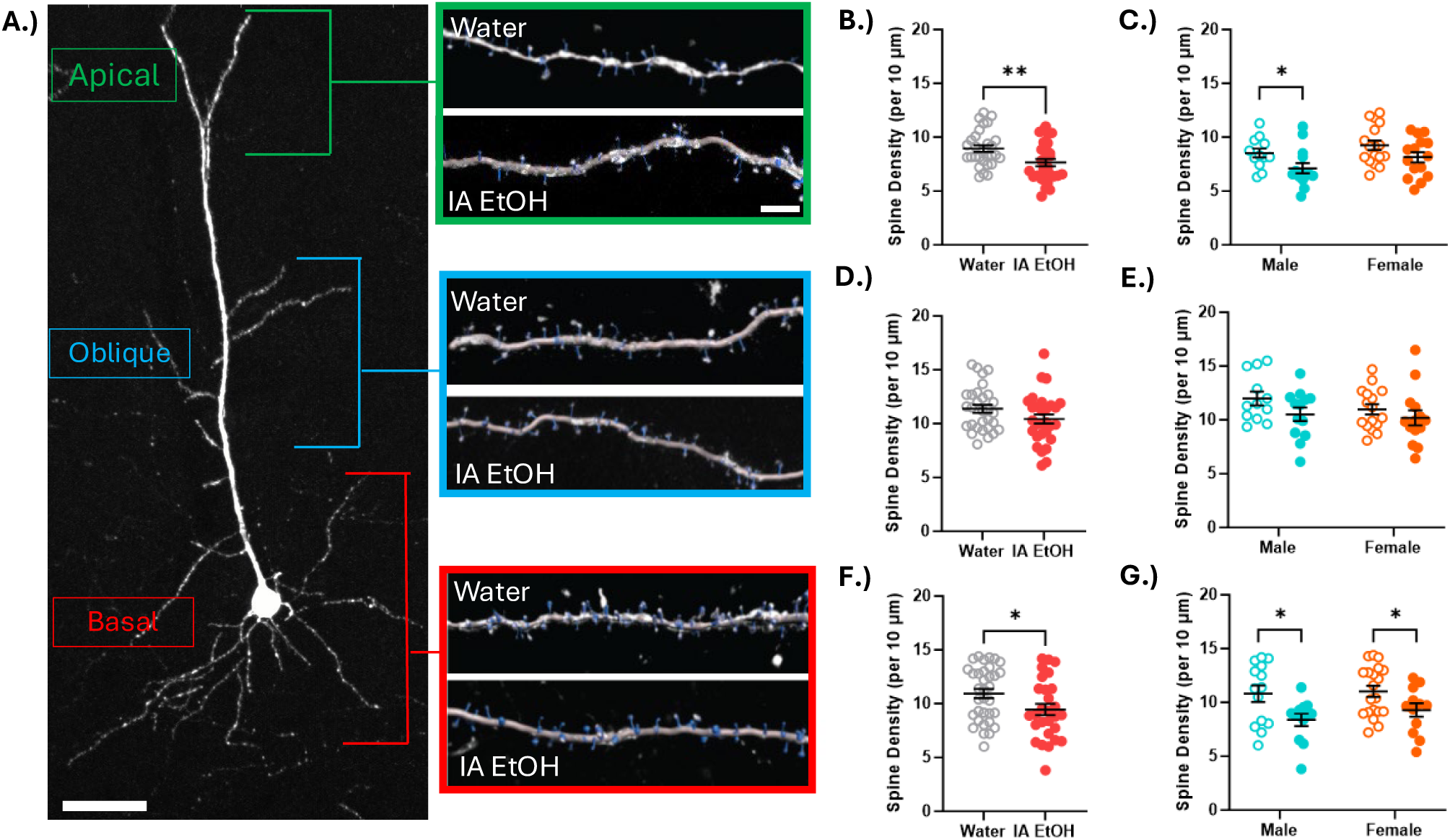
Adolescent IA EtOH reduced apical and basal spine density in PFC^MdT^ neurons: **A.)** Image of PFC^MdT^ pyramidal neuron (PN) with segments demarcated for areas of apical, oblique, and basal dendrite classification used in spine density analyses. Green (apical), blue (oblique), and red (basal) boxes show representative segments from adolescent water and IA EtOH mice with spines identified in turquoise. **B.)** Apical spine density in PFC^MdT^ PNs was reduced in apical dendrites and when separated by sex **C.**), males showed a specific effect, while females were near significant, p = 0.07. **D.)** Oblique spine density in PFC^MdT^ PNs were not significantly affected by adolescent IA EtOH, **F.)** with no differences observed when separated by sex. **F.)** Basal dendrite spine density was significantly reduced by adolescent IA EtOH, **G.)** which occurred in both male and female mice. ** p < 0.01, * p < 0.05.

Image analyses were performed using IMARIS 10.2 software where background was removed, and each segment was semiautomatically traced using the IMARIS filament function module with spine density calculated as number of spines per 10 μm. Two to four segments from separate PFC^MdT^ neurons were traced for each type of dendrite from each mouse with a confirmed injection target. Male (n = 12 mice: water = 6, IA EtOH = 6) and female (n = 14 mice, water = 8, IA EtOH = 7) mice were analyzed across 3 dendritic segments.

### Statistical Analyses

Graphing and statistical analyses were performed on GraphPad Prism 10. Alcohol dose consumed and preference between male and female mice in each group were compared using mixed-effects analysis followed by Tukey’s multiple comparisons test. Comparisons of water and IA EtOH on behavioral tasks and spine density were performed using unpaired student’s t-tests. To determine if there were sex effects on behavioral measures and spine density, sex was included as an independent variable and analyzed as a Two-Way ANOVA followed by a post-hoc comparison using Fisher’s LSD multiple comparisons test. Statistical significance was set at p < 0.05 for all comparisons.

## Results

### Intermittent Access to Alcohol (IA EtOH) during Adolescence

Male and female mice had access to 15% (v/v) ethanol and water every other day, consuming an initial dose of 6.5 g/kg on drinking session one, with male mice escalating to ∼15 g/kg by day 4 and females escalating their alcohol consumption to ∼20 g/kg on day 5 (**Figure 1C**). Analysis of alcohol dose consumed by male and female mice using a mixed-effects analysis indicated a main effect of session, F(6.087, 92.61) = 4.33, p = 0.006, and a main effect of sex, F(1,16) = 7.16, p = 0.017 with no sex by session interaction, F(14, 213) = 0.99, p = 0.46. Tukey’s multiple comparisons test found a significant difference between dose consumed by male and female mice on sessions 4, 5, 9, and 11. Preference for the alcohol bottle also escalated from ∼30% during the first session, reaching 74% in female mice by day 5, but with a more gradual increase in male mice reaching 65% on the last session (**Figure 1D**). Mixed-effects analysis of alcohol preference showed a main effect of session, F(5.173, 78.33) = 7.605, no main effect of sex, F(1,16) = 3.27, p = 0.09, and a significant sex-by-session interaction, F(14,212) = 2.36, p = 0.005. Tukey’s multiple comparisons test found a significant difference in alcohol preference between male and female mice on sessions 2, 3, and 5. On session 13, at ∼6 hours after the onset of the dark cycle and presentation of the alcohol bottles, blood alcohol concentrations demonstrated a large range of values (4-150 mg/dl) (**Figure 1E**). BAC values (mean±SEM) for males = 78.5±27.6 mg/dl and females = 59.6±27.6 mg/dl overlapped or approached the binge level threshold respectfully. A student’s t-test revealed that male and female BAC did not differ, t(df)=0.69, p = 0.5 at this timepoint, however, only male mice had a significant correlation between BACs and alcohol consumption, R^2^ = 0.845, p = 0.0095 with females at R^2^ = 0.249, p = 0.143 (**Figure 1E**).

### Adolescent IA EtOH did not affect anxiety as measured by the elevated plus maze

Chronic alcohol consumption has previously been associated with elevated anxiety levels during the withdrawal period, which could influence performance on cognitive tasks. The EPM leverages rodents’ unconditioned fear of heights and open spaces to assess anxiety; decreased entries into and/or decreased time spent on open arms reflects anxiety-like behavior ^25^. In pooled EPM data from males and females, control (n=18) and IA EtOH mice (n=17) were compared using a student’s t-test and no significant differences were found in the percent time spent on the open arms (t(33) = 0.683, p = 0.5) (**Figure 2B**) or percent open arm entries (t(33) = 0.55, p = 0.587). When data from male and female mice were compared using a 2-way ANOVA, there was no effect of sex, F(1,31) = 0.138, p = 0.713, alcohol, F(1,31) = 0.344, p = 0.562, or a sex by alcohol interaction, F(1,31) = 0.215, p = 0.647 (**Figure 2C**). For open arm entries, there were no main effects of sex, F(1,31) = 1.73, p = 0.198, or alcohol, F(1,31) = 0.438, p = 0.513, and no sex by alcohol interaction, F(1,31) = 0.002, p = 0.969. The EPM task demonstrated that the adolescent IA EtOH procedure did not cause anxiety-like behavior compared to controls at 24-32 hours post alcohol consumption in male and female mice.

### Adolescent IA EtOH did not affect spontaneous locomotor activity or anxiety on the open field task

The open field task measures spontaneous locomotor activity and provides an additional measure of anxiety-like behavior, two factors that can impact subsequent cognitive task performance (**Figure 2D, G**). Thigmotaxis, or the tendency of rodents to remain close to walls, is associated with increased anxiety and conversely, percent time spent in the center of the arena decreases with higher anxiety levels ^26^ (**Figure 2G**). For spontaneous locomotor activity, a student’s t-test revealed that there was no significant difference between IA EtOH mice (n = 18), and controls (n =18) in distance traveled, t(34) = 1.33, p = 0.193, (**Figure 2E)**. Additionally, there was no effect on percent time in the center (t(34) = 0.192, p = 0.849) (**Figure 2H**). When sex was compared using a 2-way ANOVA, we observed a main effect of sex, F(1,32) = 6.29, p = 0.017, no effect of alcohol, F(1,32) = 2.43, p = 0.129, and no sex by alcohol interaction, F(1,32) = 0.054, p = 0.818 (**Figure 2 F, I**). While male mice appeared to express less anxiety on the OFT, adolescent IA EtOH did not affect locomotor activity or anxiety-like behavior.

Results from EPM and OFT indicate that any differences observed from subsequent tasks were unlikely to be influenced by any effects of adolescent IA EtOH on motor function or anxiety.

### Adolescent IA EtOH did not affect Novel Object Recognition Task performance

The novel object recognition task (NORT) is a neophiliac task based on rodents’ tendency to explore novel stimuli. Accordingly, a mouse with intact single-item recognition memory would typically be expected to spend more time exploring the novel object than the sample object during the test phase. Following adolescent IA EtOH, a student’s t-test demonstrated that there was no significant difference in discrimination ratio between IA EtOH mice (n=17) and control mice (n=18) (t(33) = 0.154, p = 0.878) (**Figure 3B**). Importantly, both IA EtOH and control mice showed significant discrimination between novel and familiar objects, as compared to a discrimination ratio of 0.0 (IA EtOH: t(16) = 4.582, p = 0.0003; control: t(17) = 5.172, p <0.0001). IA EtOH and control groups did not differ in other aspects of performance on the NORT, including the total duration of interactions in the sample phase (t(32) = 0.8991, p = 0.3753) or test phase (t(33) = 0.635, p = 0.53), percentage of time in the center of the arena during the sample phase (t(33) = 0.734, p = 0.468), test phase (t(34) = 0.223, p = 0.825), or total distance traveled during the sample (t(32) = 1.460, p = 0.154) or test phase (t(32) = 1.749, p = 0.09). When sex was included as an independent variable, a two-way ANOVA revealed no significant main effect of sex on discrimination ratio, F(1,31) = 0.403, p = 0.53, no main effect of alcohol, F(1,31) = 0.055, p = 0.816, with no sex by alcohol interaction, F(1,31) = 1.125, p = 0.297 (**Figure 3C**). These data demonstrate that NORT performance was not affected by adolescent IA EtOH or between male and female mice.

### Adolescent IA EtOH reduced performance on the Object-in-Place Recognition Task

The OPRT assesses the ability of mice to associate objects with cued spatial locations in the arena and attend when objects have been moved from their previous configuration. IA EtOH mice (n=18) performed significantly worse on the OPRT compared to control mice (n=15) as indicated by a difference in discrimination ratio (t(31) = 2.454, p = 0.02) (**Figure 3E**). Additionally, when compared to chance interactions, control mice significantly discriminated between the objects in new and familiar locations, but IA EtOH mice did not (control: t(14) = 3.468, p = 0.004; IA EtOH: t(17) = 0.338, p = 0.739). IA EtOH and control groups did not differ in other aspects of performance on the OPRT, including the total duration of interactions in the sample (t(31) = 0.913, p = 0.368) or test phase (t(31) = 0.32, p = 0.751), percentage of time in the center of the arena during the sample (t(32) = 0.604, p = 0.55) or test phase (t(32) = 0.0805, p = 0.936), or total distance traveled during the sample (t(32) = 1.732, p = 0.093) or test phase (t(32) = 0.258, p = 0.798) (data not shown). When sex was analyzed as an independent variable, a two-way ANOVA found no significant main effect of sex on discrimination ratio F(1,31) = 1.18, p = 0.29, a main effect of alcohol, F(1,29) = 5.45, p = 0.027, and no sex by alcohol interaction F(1,29) = 0.008, p = 0.927 (**Figure 3F**). Our results demonstrate that adolescent IA EtOH impairs performance on the OPRT that was not sex-dependent.

### Spontaneous alternation on the Y-maze was not affected by adolescent IA EtOH

The Y-maze uses the natural exploratory behavior of mice to assess short-term spatial working memory, which is dependent on the hippocampus and prefrontal cortex^27–29^. Mice have an innate tendency to explore novel areas, thus, mice with intact spatial memory would be expected to enter a less recently visited arm of the maze ^28^. This behavior is described as spontaneous alternation, and a mouse with a high percentage of correct alternations is thought to have intact spatial working memory. There was no significant difference in the percentage of correct alternations between the IA EtOH (n=17) and control (n=17) groups (t(32) = 1.805, p = 0.081) (**Figure 3H**). There were no statistically significant differences observed in overall locomotor activity in the Y-maze, as measured by total number of arm entries (t(32) = 2.001, p = 0.054) and total distance traveled (t(32) = 1.87, p = 0.071). These near significant effects on performance and locomotor activity indicated that IA EtOH mice trended towards increased performance, but reduced activity on the Y-Maze. When sex was included as an independent variable and analyzed using a 2-way ANOVA, we found no effect of sex, F(1,30) = 0.176, p =0.68, or alcohol, F(1, 30) = 2.73, p = 0.109, and no alcohol by sex interaction, F(1, 30) = 0.354, p = 0.56) (**Figure 3I**). Results from the Y-maze task demonstrated that adolescent IA EtOH does not affect spatial working memory when assessed by spontaneous alteration.

### Reduced spine density in PFC^MdT^ neurons following adolescent IA EtOH

To determine if PFC^MdT^ neurons underwent synaptic alterations following adolescent IA EtOH, virally transduced layer V pyramidal GFP positive neurons were imaged from behavioral mice. Apical, oblique, and basal dendrite segments were specifically targeted for analyses. For apical segments, statistical analyses using a student’s unpaired t-test revealed a significant reduction of adolescent IA EtOH on spine density, t(56) = 2.834, p = 0.006 (**Figure 4B**). When sex was analyzed as an independent variable, we did not observe a main effect of sex on apical spine density, F(1,54) = 3.66, p = 0.061, but the main effect of alcohol remained, F(1,54) = 7.6, p = 0.008, with no sex by alcohol interaction, F(1,54)=0.102, p = 0.751 (**Figure 4C**). When uncorrected Fisher’s LSD post hoc comparisons were applied, IA EtOH males had a significant reduction in apical spine density, p = 0.04, while IA EtOH females did not reach significance, p = 0.07. The spine density for oblique segments was not statistically significant, unpaired t-test t(57) = 1.64, p = 0.106 (**Figure D**). Additionally, oblique segments had no main effect of sex, F(1.51)=1.21, p = 0.278, no main effect of alcohol, F(1,51) = 3.409, p = 0.071, and no sex by alcohol interaction, F(1,51) = 0.296, p = 0.589 (**Figure 4E**). Finally, we found a significant decrease in basal spine density, t(60) = 3.39, p = 0.03, following adolescent IA EtOH. When sex differences for basal dendrites were analyzed, a pattern similar to apical dendrites was observed with sex having no main effect, F(1,54)=0.806, p = 0.37, a significant main effect of alcohol, F(1.54) = 10.95, p = 0.0017, and no sex by alcohol interaction, F(1,54) = 0.297, p = 0.59 (**Figure 4 F,G**). Yet, multiple comparisons showed both males, p = 0.011, and females, p = 0.047 reached significance for reductions in basal spine density between water and IA EtOH. These results demonstrate that IA EtOH reduces basal and apical spine density largely independent of sex.

## Discussion

The present study sought to identify behavioral deficits in a relatively rapid battery of cognitive assessment tasks following adolescent alcohol consumption in male and female mice. We hypothesized that PFC^MdT^ projections were vulnerable to adolescent IA EtOH with an associated behavioral phenotype consistent with PFC^MdT^ dysfunction. The main finding from our behavioral measures is that adolescent IA EtOH results in selective deficits in spatial object recognition on the OPRT in both male and female mice, without affecting locomotion, anxiety, novel object recognition, or Y-maze spontaneous alternation. We additionally assessed spine density of PFC^MdT^ neurons to link behavioral findings with structural changes to PFC^MdT^ circuitry. Spine density measurements of PFC^MdT^ neurons from mice following the behavioral assessment indicated that IA EtOH led to a reduction in spine density in apical and basal, but not oblique dendritic segments, independent of sex.

Key advantages of the 2-bottle choice IA EtOH procedure are that mice voluntarily consume alcohol in a goal-directed manner under low stress conditions, preserving oral ingestion and alcohol pharmacodynamics, akin to human alcohol consumption. Under the IA EtOH schedule, mice demonstrated an escalation of alcohol consumption during adolescence, higher preference for alcohol, and a subset showed blood alcohol concentrations reaching binge levels, a pattern consistent with previous reports ^6, 30^. In comparison with values from previous studies using the IA EtOH, adolescent male mice in this study consumed less alcohol per day (mean range: ∼6 to 15 g/kg vs. ∼9 to 18 g/kg^30^ and 10 to 18 g/kg^6^), had a noticeably lower preference for the alcohol bottle (mean range: 30 to 60% vs. 50-90%^30^ and 40 to 85%^6^). These observed differences could have been influenced by diet as autoclaved Teklad mouse chow used in this study (TL2019S) is known to reduce alcohol consumption, preference, and BACs when directly compared to LabDiet 5001 used in previous IA EtOH studies ^6, 31^. To assess the range of blood alcohol levels, samples were taken 6 hours into the drinking session, providing BAC values at a single time point and unlikely to reflect peak BACs. We did observe a significant correlation between dose consumed and BACs for male mice which could either indicate higher consumption nearer blood sampling in males, higher alcohol metabolism or titration with water in females. Voluntary consumption procedures with more limited alcohol access periods like drinking in the dark (DID) paradigms would more likely capture peak BACs in both sexes ^32^, and while forced alcohol exposure can obviate variability in BACs, the tradeoff is a lack of volitional intake and associated alcohol pharmacokinetics. A direct comparison of voluntary and involuntary adolescent exposure methods on PFC^MdT^ relevant neurobiological and behavioral consequences is warranted. Considering these shortcomings of the procedure, we observed a selective behavioral effect and structural neuroadaptations in the adolescent IA EtOH procedure, further validating its continued use.

Behavioral testing began one day after the cessation of alcohol access, so it was important to assess presence of anxiety-like behavior and locomotor activity to absolve any significant withdrawal effects that could impact subsequent tasks focused on cognition. Adolescent IA EtOH mice did not demonstrate significant differences in anxiety as assessed by EPM and OFT center time and we did not detect any differences in locomotor activity across all 5 tasks. Previous studies using adolescent IA EtOH in male mice have also not observed effects on the EPM at 2 or 72 hours or handling-induced convulsions at 2 and 8 hours following drinking cessation^6, 33^. Collectively, the adolescent IA EtOH consumption paradigm consistently does not result in observable somatic withdrawal symptoms in males or withdrawal-related anxiety in males and females as might be observed in forced exposure methods or more long-term IA EtOH access^34^. We did find that female mice showed more anxiety-like behavior in the OFT, which has not typically been reported^35^. A caveat of the present study and others that require measuring individual drinking values on the 2-bottle choice task is that all mice were single-housed throughout the adolescent period and so social isolation may be a contributing factor to basal anxiety differences between male and females not previously observed. In our study, water control mice were also single-housed and although we have previously shown that housing conditions during adolescence do not affect intrinsic electrophysiological properties of medial PFC neurons^6^, single-housed mice are known to have increased dendritic spine density of immature thin spines in the mPFC compared to group-housed mice^36^ which may make them more vulnerable to adolescent IA EtOH, a potential interaction of housing conditions and alcohol.

Experimental findings from our memory-based tasks demonstrate that adolescent IA EtOH have a selective impairment on spatial object recognition, but not in the novel object recognition task. These findings are consistent with deficits in allocentric versus egocentric processing, where single object recognition is intact, however associative recognition memory is impaired. In the NORT, both groups and sexes showed a statistical preference for the novel object in the test phase, but in the OPRT, only control male and female mice showed discrimination of switched object locations. Importantly, there were no significant differences observed in the total duration and number of interactions with objects in the NORT or OPRT and no differences in locomotor activity or anxiety-like behavior measured by center time during the task. Therefore, OPRT deficits are likely the result of the learning and memory task requirements. Several studies have assessed object recognition performance following passive adolescent alcohol exposure methods including alcohol injections, alcohol gavage, and alcohol vapor. Adolescent alcohol gavage in rats has resulted in deficits in NORT^20^ and long-term effects in female rats. Adolescent, but not adult alcohol gavage in mice ^21^ impairs NORT performance and adolescent intraperitoneal alcohol injections impair object recognition and sensitivity to the temporal but not spatial processing of object location^22^. In a similar 2-bottle choice alcohol consumption paradigm to the present study, but initiated following weaning (PND 21-52), alcohol drinking mice were not impaired in NORT, but did show impairment on object location memory that persisted into abstinent adulthood^37^. In contrast to our data, they observed a significant anxiety, locomotor, and withdrawal phenotype post alcohol that may have influenced spatial memory performance including Y-maze deficits and perhaps, a consequence of an earlier initiation of alcohol consumption. In summary, previous studies and the results here demonstrate that several adolescent alcohol exposure models lead to impairment on object recognition tasks, with higher alcohol passive exposures leading to deficits on object recognition, and voluntary consumption models more likely to affect associative memory processes involved in recognizing object location.

Importantly, studies have been performed that can attribute aspects of object recognition to discrete brain regions and disassociate their respective roles. Hippocampal and perirhinal lesions disrupt performance on the NORT and our lack of effect on this task in addition to a lack of anxiety effects suggests limbic brain structures were not severely affected by adolescent alcohol. Our finding is more consistent with mPFC and MdT dysfunction as bilateral lesions of mPFC or MdT selectively impair OPRT, but not NORT performance^38^. Furthermore, disconnection of the MdT and mPFC tracts additionally impaired OPRT without affecting single-item recognition memory on the NORT, providing further evidence that the PFC and MdT bilateral circuitry are critical to associative recognition memory^38^. We propose that the deficits seen in IA EtOH mice on the mPFC- and MdT-dependent OPRT, but not on other behavioral tasks, selectively disrupt the mPFC^MdT^ circuit and that this represents a disassociation from hippocampal and parahippocampal effects of adolescent alcohol, however, further interrogation using circuit-based techniques and *in vivo* monitoring of circuit activity during task performance are needed to fine tune this conclusion.

Adolescent IA EtOH did not affect spontaneous alternation on the Y-maze. On the contrary, there was a nearly significant increase in performance in IA EtOH mice and decrease in total arm entries. Performance on the Y-maze has been shown to be dependent on the integrity of the hippocampus and, in some cases, the PFC ^27, 28^. A previous study found that ethanol i.p. injections in rats during adolescence impaired spontaneous alternation on the Y-maze ^39^, yet female mice performance was not affected using alcohol fed mice using the Leiber Di Carli diet ^40^. In the present study, differences in findings may be a result of different models of alcohol consumption (2.5g/kg ethanol intraperitoneal injection versus voluntary IA EtOH). Detection of spatial working memory deficits of adolescent alcohol may require more challenging working memory conditions such as the delayed non-match to sample T-Maze, where reduced performance has been observed in male mice following adolescent IA EtOH^6^.

In male and female mice, we demonstrated that adolescent IA EtOH reduced spine density on basal and apical dendrites of layer V PFC^MdT^ pyramidal neurons (PNs), consistent with a dampening of synaptic activity. PFC^MdT^ PNs in layer V are typically grouped within the pyramidal tract categorization of PN subtypes within the mPFC, and recent neuroanatomical and functional studies have begun to map out the local synaptic organization of mPFC PN subtypes^41^. For instance, the tufted collection of apical dendrites from pyramidal tract PNs in the mPFC are biased towards integrated excitatory inputs from the ventromedial thalamus and the MdT, related to sensorimotor processing and rule strategy^16, 42^. Their basal dendrites receive primarily inputs from local mPFC PNs but also receive long-range inputs from the basal lateral amygdala and ventral hippocampus^41^. While the synaptic organization of the PFC is not fully resolved, our morphological data would suggest a weakening of synaptic inputs on PFC^MdT^ PNs carrying sensory, associative, and contextual information from thalamic and limbic structures.

Given that the MdT links hippocampal and prefrontal networks supporting relational memory and object-in-place recognition ^12, 13, 43^, reduced connectivity between the mPFC and MdT may specifically impair binding object features to spatial context while sparing novelty detection.

These circuit-level alterations are likely a consequence of IA EtOH pharmacological actions on normal synaptic pruning during adolescence. The medial PFC undergoes a developmental surge in spine formation followed by activity-dependent elimination and stabilization, accompanied by high synaptic turnover during adolescence ^44, 45^. Alcohol exposure, known to acutely enhance GABAergic and reduce glutamatergic activity^46, 47^, during this window could accelerate synaptic pruning or destabilize maturing PFC^MdT^ synapses leading to persistent loss of thalamic feedback and reduced excitatory tone as supported by our spine density results. Such developmental disruption provides a mechanistic basis for selective deficits in object-in-place recognition after adolescent drinking and highlights the potential vulnerability of the adolescent PFC to alcohol-induced remodeling of cortico-thalamic circuits.

In summary, mice that consumed alcohol during adolescence had deficits in object-in-place associative recognition memory but not in single-item novel object recognition consistent with behavioral performance following lesions of the medial PFC or MdT subregions. The results from this study imply that the connectivity between the mPFC and MdT may be a locus of adolescent alcohol’s effects on cognition further evidenced by a reduction in the spine density of PFC^MdT^ projections. This circuit has been implicated in cognitive control and if it is similarly affected in humans is likely to contribute to the challenges in treating alcohol use disorder in individuals with histories of excessive alcohol use during adolescence including relapse vulnerability and adherence to treatment such as cognitive behavioral therapy.

## Acknowledgements

This work is funded by grants AA030619 and AA024507. The authors declare no conflict of interest.

---

Figure 1A created in BioRender. Salling, M. (2025) https://BioRender.com/vvrapcu

## Notes

### Competing Interest Statement

The authors have declared no competing interest.

